# Simultaneous tracking of *Pseudomonas aeruginosa* motility in liquid and at the solid-liquid interface reveals differential roles for the flagellar stators

**DOI:** 10.1101/541615

**Authors:** Andrew L. Hook, James L. Flewellen, Jean-Frédéric Dubern, Alessandro Carabelli, Irwin M. Zaid, Richard M. Berry, Ricky Wildman, Noah Russell, Paul Williams, Morgan R. Alexander

## Abstract

Bacteria sense chemicals, surfaces and other cells and move toward some and away from others. Studying how single bacterial cells in a population move requires sophisticated tracking and imaging techniques. We have established quantitative methodology for label-free imaging and tracking of individual bacterial cells simultaneously within the bulk liquid and at solid-liquid interfaces by utilizing the imaging modes of digital holographic microscopy (DHM) in 3D, differential interference contrast (DIC) and total internal reflectance microscopy (TIRM) in 2D combined with analysis protocols employing bespoke software. To exemplify and validate this methodology, we investigated the swimming behavior of *Pseudomonas aeruginosa* wild type and isogenic flagellar stator mutants (*motAB* and *motCD*) respectively within the bulk liquid and at the surface at the single cell and population levels. Multiple motile behaviours were observed that could be differentiated by speed and directionality. Both stator mutants swam slower and were unable to adjust to the near surface environment as effectively as the wildtype highlighting differential roles for the stators in adapting to near surface environments. A significant reduction in run speed was observed for the *P. aeruginosa mot* mutants, which decreased further on entering the near-surface environment. These results are consistent with the *mot* stators playing key roles in responding to the near-surface environment.

**Importance:** We have established a methodology to enable the movement of individual bacterial cells to be followed within a 3D space without requiring any labelling. Such an approach is important to observe and understand how bacteria interact with surfaces and form biofilm. We investigated the swimming behavior of *Pseudomonas aeruginosa*, which has two flagellar stators that drive its swimming motion. Mutants that only had either one of the two stators swam slower and were unable to adjust to the near surface environment as effectively as the wildtype. These results are consistent with the *mot* stators playing key roles in responding to the near-surface environment, and could be used by bacteria to sense when it is near a surface.

## Introduction

Flagella and type IV pili (TFP) mediated motility enable bacteria to migrate towards nutrients or away from toxic substances (5, 49) and play key roles in bacterial biofilm formation and host-pathogen interactions (25, 46). In addition, these bacterial appendages also play a role in surface sensing (30). The mechanisms by which individual cell behaviors drive social phenomena such as swarming, twitching and biofilm development have been the subject of intense investigations (1, 13–14, 16, 23–25, 29, 31, 34, 40, 47). The ability to collect data simultaneously on individual bacterial cells in a population that includes both planktonic and surface attached cells will facilitate the acquisition of spatio-temporal dynamics that will aid our understanding of the signal transduction mechanisms that drive bacterial social behaviors.

*Pseudomonas aeruginosa* is a Gram-negative rod-shaped cell with a single polar flagellum that employs a number of different mechanisms for moving through liquids and across surfaces. These include flagella-mediated swimming, spinning, near-surface swimming and swarming, type IV pili-mediated twitching (crawling and walking), gliding (active movement without the use of flagella or pili), and sliding i.e. passive movements over surfaces through the use of surfactants (14, 22). Flagella are also associated with bacterial surface mechanosensing and play a key role during the early stages of biofilm formation (3, 24, 29, 34, 39). In *P. aeruginosa*, the flagellum, in contrast to most other bacterial species, is driven by two (rather than one) stator complexes termed MotAB and MotCD. These are the static elements of the bacterial motor that generate torque for flagellar rotation powered by proton-motive force (2). Either MotAB or MotCD is sufficient for swimming but the two stators make differential contributions to swarming motility and biofilm formation (15, 39, 40). Deletion of *motCD*, but not *motAB*, renders *P. aeruginosa* incapable of swarming over agar at concentrations above 0.5 % (40), suggesting that the MotCD stator complex is able to apply torque more efficiently than the MotAB complex (15, 24, 40). Both stators are required for biofilm formation (39).

Recent studies suggest an interaction between MotCD and the second messenger cyclic diguanylate (c-di-GMP), which plays a key role in driving the lifestyle switch of motile *P. aeruginosa* cells towards surface-associated biofilm formation (42), through the PilZ domain-containing protein FlgZ (1). This interaction may cause the MotCD stator to be displaced from the flagella motor or may increase the likelihood of this event occurring (24). Moreover, free MotC is able to interact with the c-di-GMP receptor, SadC, causing it to become activated triggering further production of c-di-GMP (3). This produces a positive feedback loop mediated by the flagella machinery that directs cells from the motile to sessile state with stator exchange playing a key signaling role. MotAB is also necessary for triggering type IV pili formation after cell attachment, further implicating a role for both stators in surface sensing (25). The contribution of flagella stators as surface sensors for biofilm formation which involves increasing production of c-di-GMP and pili, has also been observed for the single stator species *Caulobacter cresentus* (16, 21). Understanding the respective roles of MotAB and MotCD in the initial interactions of swimming cells with surfaces is important particularly in the context of early stage biofilm formation.

Cell tracking in 3D allows for a complete analysis of bacterial movement and can be used to assess the trajectory of a bacterium approaching or leaving a surface. Early work following single cells determined the z-position by tracking the focal plane of a bacterium and recording x and y movements optically. This method can only follow a single cell at a time and is thus unable to record the movement of many cells in a population (18). 3D volumes have been imaged using optical techniques with narrow focal planes such as confocal microscopy to acquire z-stacks, but this approach is limited by low light transmission and consequently low acquisition rates. Alternatively, ‘lookup’ tables have been used to match the image of an object with reference images taken at known distances away from the focal plane of an object (27, 36, 47).

Most recently, 3D tracking of bacteria has been achieved using digital holographic microscopy (DHM) where the x, y, z-position of individual cells is captured via recorded diffraction patterns and determined computationally (17, 44). For the monotrichous *P. aeruginosa,* swimming in liquids has been analyzed using DHM where motility patterns have been classified as *oscillation*, *meander*, *helix*, *pseudohelix* and *twisting* (44). The key advantage of DHM is its ability to observe the 3D trajectories of many cells in a bacterial population simultaneously; however, features such as their orientation and shape are difficult to ascertain from the holograms. A recent variant using a three laser setup permitted the use of DHM for determining bacterial volume and orientation throughout a 3D space. However, this approach has thus far only been demonstrated for the analysis of single bacterial cells released near a surface using optical tweezers (8).

To gain novel insights into the differences in cell behavior within the bulk and at the surface, it is necessary to image both environments concurrently. To achieve this, we developed a novel multimode 2/3D microscope with interlaced acquisition of 2D surface differential interference contrast (DIC) or total internal reflectance microscopy (TIRM) images with 3D DHM. The spatial and temporal resolution of the DIC images at the surface were sufficient to allow the bacterial cell orientation to be determined, whilst TIRM imaging confirmed the close proximity to the surface (<200 nm) of near surface swimming cells. In addition, to enable imaging of a higher density bacterial population than is possible for DHM, we incorporated an optical relay (10) to recreate the sampling area virtually, enabling it to be rapidly scanned in the z-dimension using a remote piezo-driven objective to avoid sample disturbance. A schematic of the multimode 2/3D microscope setup is shown in Fig. 1A-B. Using this methodology we were able to simultaneously determine the individual cell orientation and proximity to the surface as well as the 3D location of cells at low and high densities over time. To exemplify the utility of this system, the microscope was used to explore the respective contributions of the *P. aeruginosa* MotAB and MotCD stators to bulk liquid movement and interactions at the solid-liquid interface.

**FIG. 1.**
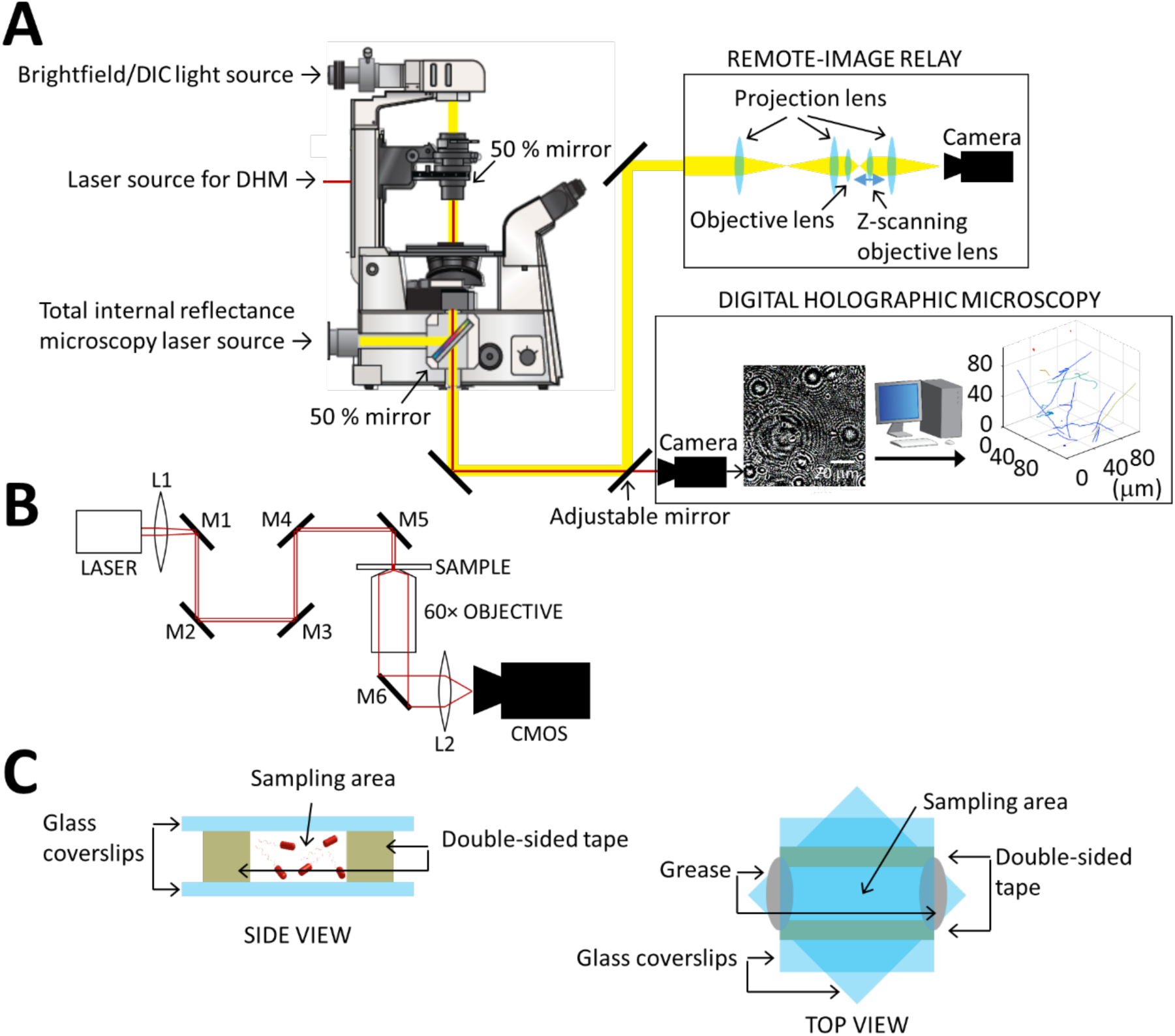
Overview of the experimental setup. (A) Schematic of the multimode 2/3D microscope used in this study including the optical ensemble for the remote-image relay. The Remote Image Relay allows scanning of the focal plane to reconstruct z-stacks without disturbing the sample. The microscope is also equipped with lasers to enable digital holographic microscopy (DHM) and total internal reflectance microscopy (TIRM). (B) A schematic diagram of the light path used for inline DHM. The positions of L1, M1 and M2 were adjustable to alter the position and focus of the laser. (B) Schematic of the sample format used in this study including a side and top view. The z-height of the sample volume (distance between glass slides) was 100 μm. The typical separation between the two strips of double-sided tape was 5 mm. The two coverslips were orientated at 45° to each other to enable the easy sealing of the channel ends with hydrocarbon grease.

## Materials and Methods

### Bacterial strains and growth conditions

*P. aeruginosa* strain PAO1 (Washington sub-line) and the isogenic *motAB*, *motCD* and *motABCD* mutants were each grown at 37°C overnight in lysogeny broth (LB) and diluted to an OD_600_ 0.01 in LB prior to incubating for a further 4 h at 37°C with shaking to reach log phase at an OD_600_ ≈ 1. Bacterial cultures were diluted to OD_600_ 0.015 in LB before inoculation into analysis chambers. The *mot* mutants were constructed by two-step allelic exchange as previously described (28). Two PCR products amplifying the upstream and the downstream nucleotide regions of *motAB* or *motCD* were generated using the primer pair 1FW/1RW, and 2FW/2RW, respectively (Table S1). Both PCR products were fused by overlapping PCR to create a deletion in the corresponding gene. The resulting fragment was cloned into the suicide plasmid pME3087 (45). Following transformation into the target strain by conjugation, single cross-overs were selected on tetracycline (125 μg.ml^-1^). The double crossover mutants were selected by carbenicillin enrichment (28). After 3 rounds of counter-selection, the resulting *P. aeruginosa* colonies were screened for the loss of antibiotic resistance by plating on LB agar supplemented with or without tetracycline. The in-frame deletions were confirmed by PCR and DNA sequence analysis.

**TABLE 1.** Summary of quantification of surface tracks including both surface and bulk measurements. Speeds presented are the average speeds measured over frame intervals of 24 ms. Speed variance is the average standard deviation for the instantaneous speeds measured for an entire trajectory. The ranges quoted are one standard deviation of such measurements for each cell. WT = wildtype.

|  | *Surface* |  |  | *Bulk* |  |  |
| --- | --- | --- | --- | --- | --- | --- |
|  | WT | *motAB* | *motCD* | WT | *motAB* | *motCD* |
| **Classification of cells** |  |  |  |  |  |  |
| Number of tracks | 150 | 138 | 142 | 71 | 154 | 97 |
| Motile (%) | 27 ± 6 | 77 ± 9 | 77 ± 20 | 97 ± 7 | 97 ± 3 | 100 ± 0 |
| -runs (% of motile cells) | 51 ± 6 | 10 ± 2 | 25 ± 28 | 64 ± 31 | 45 ± 10 | 46 ± 34 |
| -near surface runs (% of motile cells) | 18 ± 9 | 54 ± 4 | 43 ± 16 |  |  |  |
| -rambling (*surface*) / oscillating (*bulk*) (% of motile cells) | 31 ± 12 | 36 ± 5 | 32 ± 21 | 36 ± 31 | 55 ± 10 | 54 ± 34 |
| Attached (%) | 72 ± 9 | 23 ± 9 | 21 ± 22 |  |  |  |
| -polar attached (% of attached cells) | 30 ± 14 | 73 ± 26 | 55 ± 31 |  |  |  |
| -long-axis attached (% of attached cells) | 70 ± 14 | 27 ± 26 | 45 ± 31 |  |  |  |
| Floating (%) | 2 ± 3 | 0 ± 1 | 2 ± 3 | 3 ± 7 | 3 ± 3 | 0 ± 0 |
| **Direction change analysis** |  |  |  |  |  |  |
| Frequency of reversal events for cell runs (Hz) | 0.9 ± 0.4 | 0.2 ± 0.0 | 0.6 ± 0.3 | 0.5 ± 0.1 | 0.1 ± 0.0 | 0.1 ± 0.1 |
| Oscillating frequency |  |  |  | 3.1 ± 1.0 | 1.5 ± 1.0 | 2.6 ± 0.6 |
| Rambling frequency (Hz) | 3.2 ± 2.1 | 2.9 ± 1.5 | 6.1 ± 0.7 |  |  |  |
| **Arrival and departure events from the surface** |  |  |  |  |  |  |
| Attachment rate (% of non-attached cells per second) | 0.2 ± 0.4 | 0.3 ± 0.4 | 0.8 ± 0.7 |  |  |  |
| Detachment rate (% of attached cells per second) | 0.2 ± 0.3 | 2.2 ± 2.5 | 2.5 ± 1.6 |  |  |  |
| **Motility characterisation** |  |  |  |  |  |  |
|  | *Surface run* |  |  | *Run* |  |  |
| Speed (µm/s) | 55 ± 7 | 18 ± 5 | 22 ± 21 | 59 ± 4 | 29 ± 9 | 45 ± 18 |
| Speed variance | 23 ± 6 | 10 ± 6 | 16 ± 12 | 18 ± 4 | 9 ± 1 | 32 ± 36 |
| K<sub>MSD</sub> (µm<sup>2</sup>/s) | 1.9 ± 0.2 | 1.9 ± 0.3 | 1.8 ± 0.2 | 1.9 ± 0.3 | 1.8 ± 0.4 | 1.9 ± 0.2 |
|  | *Rambling* |  |  | *Oscillating* |  |  |
| Speed (µm/s) | 18 ± 8 | 13 ± 4 | 12 ± 4 | 12 ± 5 | 19 ± 5 | 10 ± 2 |
| Speed variance | 11 ± 5 | 8 ± 4 | 11 ± 3 | 11 ± 8 | 9 ± 2 | 5 ± 1.3 |
| K<sub>MSD</sub> (µm<sup>2</sup>/s) | 1.7 ± 0.3 | 1.7 ± 0.4 | 1.5 ± 0.4 | 1.2 ± 1 | 1.6 ± 0.4 | 1.3 ± 0.3 |

For genetic complementation of the in-frame deletion mutants the *motAB* and *motCD* genomic regions were PCR-amplified using *P. aeruginosa* chromosomal DNA as a template and the primer pairs indicated in Table S1. The purified fragments were cloned into the shuttle vector pME6032 (19). The insertions were verified by restriction enzyme and sequence analysis and introduced into the *motAB* and *motCD* mutants by electroporation. The swarming ability of the *mot* mutants was verified by swarming plate assays (data not shown).

### Sample Chamber

Two strips of double-sided tape, approximately 2 × 15 × 0.1 mm, were placed on a borosilicate glass coverslip (VWR) in parallel with a gap of ∼4 mm. A second glass coverslip was placed on top of the double-sided tape to create a chamber with a floor and ceiling of glass and walls of double-sided tape. The second coverslip was rotated 45° to allow ease of loading the channel with growth medium (Fig. 1C). After inoculation with bacteria, the two ends of the chamber were sealed using silicone free grease (Apiezon).

### Microscopy

Imaging was achieved using a bespoke multimode microscope (Fig. 1A; Cairn Ltd.). Samples were analyzed at 37°C using a Nikon Eclipse Ti inverted microscope using a 60×, NA=1.49, WD=0.13 mm oil objective. The microscope was fitted with an environmental chamber (Okolab) to regulate temperature, relative humidity and CO_2_. Images were acquired using an Orca-Flash 4.0 digital CMOS camera (Hamamatsu) at a typical acquisition rate of 41.6 Hz. DIC imaging was achieved using a single channel white MonoLED (Cairns) light source. A polariser was inserted above the condenser, and Wollaston prisms were inserted between the condenser and polariser and below the objective. Use of the prisms did not adversely affect DHM image quality. Inline DHM imaging was acquired using a 685 nm LX laser (Obis). TIRM was conducted using an Obis 488 nm LX laser (Cairn) controlled using an illumination system (iLas2). The sample was illuminated through the objective by use of a 50 % mirror, controlled to achieve total internal reflectance by irradiating the sample at an incidence angle (*θ*) greater than the critical angle (*θ_c_*) given by equation 1, where *n*_1_ is the refractive index of the incident medium and *n*_2_ is the refractive index of the transmission medium.

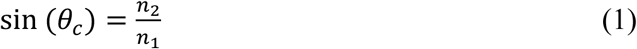

Bacterial imaging was achieved through the local change in refractive index and *θ_c_* induced upon cells entering the evanescent field caused by the TIR. The penetration depth (*d*) of the evanescent field was determined using equation 2, where *θ* is the wavelength of the light = 488 nm and tuned to 200 nm.

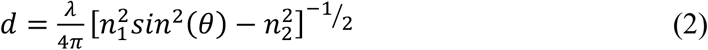

A total of 5 biological repeats for each mutant was prepared and imaged by interlaced capture of DHM with either DIC or TIRM. Each image sequence consisted of 2000 frames (1000 frames for each acquisition mode) for a total duration of 48 s.

The optical requirements of DHM, DIC and TIRM were met without interfering with the other imaging modes, thus, multimode acquisition could be achieved by modulating the intensity of the different illumination sources. Different modes of acquisition could not be acquired simultaneously but required interlaced capture achieved through computer control. The limiting factor in the time resolution of bacterial motion capture was the frame rate of the camera, which in this case was 100 Hz.

Z-stack images were acquired using a bespoke remote focusing assembly as described previously (10).

To sample the early stages of bacterial cell surface attachment, *P. aeruginosa* was inoculated into the sample chamber at OD_600_ 0.015 in LB. Bacteria were observed to undergo cell division within the chamber for 4-6 h until reaching stationary phase.

To observe flagella orientation during reversals, PAO1 was fluorescently stained using Alexa Fluor carboxylic acid succinimidyl esters (Alexa Fluor 488, ThermoFisher; Friedlander et al. 2013; Turner, Ryu, and Berg 2000). PAO1 cells were grown overnight in LB medium at 37 °C and 200 rpm, resuspended and diluted in fresh LB medium at OD_600_ = 0.01. When cells reached exponential phase, they were centrifuged at 2,000 x g for 10 min and the growth medium removed. The cell pellet was gently resuspended in a wash buffer of 10^−2^ M KPO_4_, 6.7 x 10^−2^ M NaCl, 10−4 M EDTA, pH-adjusted to 7.0 with HCl and centrifuged. After two additional rinses, cells were incubated with 0.5 mg/mL Alexa Fluor 488 carboxylic acid succinimidyl ester for 1 h at room temperature with gentle rocking. Residual dye was removed after washing twice and the cells resuspended in PBS for imaging.

### Image processing

After acquisition, holograms were processed using a bespoke Matlab script (ImageProcess). Image intensity across a stack was normalised, the median image over the stack was subtracted to remove background signal including attached cells. DHM image output sequences were reconstructed using a bespoke Python package (See-Through Scientific) (17–51) making use of Rayleigh-Sommerfeld formalism to determine bacterial X-Y-Z coordinates. Bacterial trajectories were determined using a bespoke Matlab script (DHMTracking). To exclude noise a voxel limit was set and applied to each image such that a similar number of objects were identified in each frame. Visual inspection of interferogram image sequences enabled an estimate of the number of bacteria per frame to ensure the voxel limit was set so as not to exclude bacterial cells. Bacterial trajectories were determined using the bespoke Matlab script (tracking). Objects were matched with their nearest neighbour within the next frame, applying a distance limit based upon the maximum speed of bacteria (100 μm/s). The script looked up to 4 frames ahead for a matching object. All tracks were also visually inspected using a bespoke Matlab script (Graphing) to ensure trajectories skipping more than 4 frames could be joined. All Matlab scripts used for processing the data are published as a part of the Supplementary information and the combined DHMTrack project can be accessed at GitHub (52).

The mean squared displacement (MSD) of tracks were calculated using a bespoke Matlab script (ROC_MSD) and calculated according to equation 3, where *x⃗*(*t_j_*) = *R⃗_j_* and was the vector of the j^th^ point on the trajectory and the angled brackets indicate all measurements within a given time t_i_ (41).

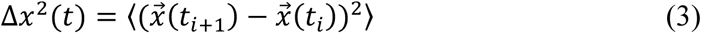

DIC images and Z-stacks were processed using a bespoke Matlab script (StackMaster). Each image was flattened by a line-by-line polynomial fit. A threshold was then applied to binarise the image to segment pixels with bacteria. Threshold was set as signal^2^:noise^2^ > 10. The square of the signal was measured so as to identify both bright and dark regions associated with the DIC image. Objects less than 20 pixels were excluded as noise. Holes within objects were filled using the Matlab function ‘imfill’. Bright and dark regions were paired based upon proximity and a common directional vector between objects within the same frame. The centre of mass of objects was calculated giving equal weighting to bright and dark regions. The length and width of the smallest rectangle around an object and the orientation of the bacterial cells was determined using a bespoke Matlab script (ParticleAnalysisDIC). For DIC Z-stacks, objects on consecutive frames were analyzed to identify overlapping pixels considering the X-Y dimensions. If common pixels were identified objects were grouped to form a single object and the centre of mass was determined for this object based upon the combined pixels using a bespoke Matlab script (DICZStack_ImageProcessing). Bacterial trajectories were determined as described for DHM data.

TIRM images were processed using ImageJ 1.50b. When processing interlaced DHM and TIRM images the centre of mass of moving cells as determined by DHM was shifted to the average position between two adjacent frames to account for the 20 ms offset between the capture of TIRM and DHM images.

### Statistical analysis

Differences between datasets were assessed using unpaired *t-*tests or one way ANOVA analysis as appropriate to determine the value *p.* The value *N* indicates the number of biological replicates. Error bars indicate ± 1 standard deviation unit.

## Results

### Imaging of individual *P. aeruginosa* cells in the bulk liquid and at the solid-liquid interface

To characterize *P. aeruginosa* motility using the multimode microscope, interlaced capture of either 2D DIC or TIRM at the surface and 3D inline DHM images in the bulk liquid respectively was conducted within 5 min of motile cells being added to the sample chamber (Fig. 1C) at 41.7 Hz (total frame rate). At this rate, bacterial fluctuating motion was under-sampled (body and Brownian motion) but directional motion was oversampled (Fig. S1). After data acquisition, bacterial trajectory generation was achieved using bespoke Matlab scripts (DHMTracking and StackMaster). Using this approach, trajectories could be captured within the bulk medium (Fig. 2A) and at the glass surface-liquid interface (Fig. 2C-D). Fig. 2B illustrates the DIC Z-stack of bacterial cell 3D positions at inoculation and after 4 h growth using the remote image relay to demonstrate that single cells in dense populations can be located and tracked. Using DIC imaging at the surface, the orientation of the bacterial cells could also be determined to establish whether a cell was orientated in the direction of travel and if cells were attached via their long axes or poles (Fig. 2C). TIRM allowed the observation of cells within a close proximity to the surface since objects were only detected within the depth of penetration of the evanescent wave into the bulk liquid (200 nm in the configuration used; Fig. 3A). The trajectories of the cells in a bacterial population were observed through the interlaced capture of DHM and DIC images. The resulting tracks were combined and where bacteria approached the surface information from both the DHM and DIC images was included. An example of a track travelling from the bulk to the surface is shown in Fig 2D.

**FIG. 2.**
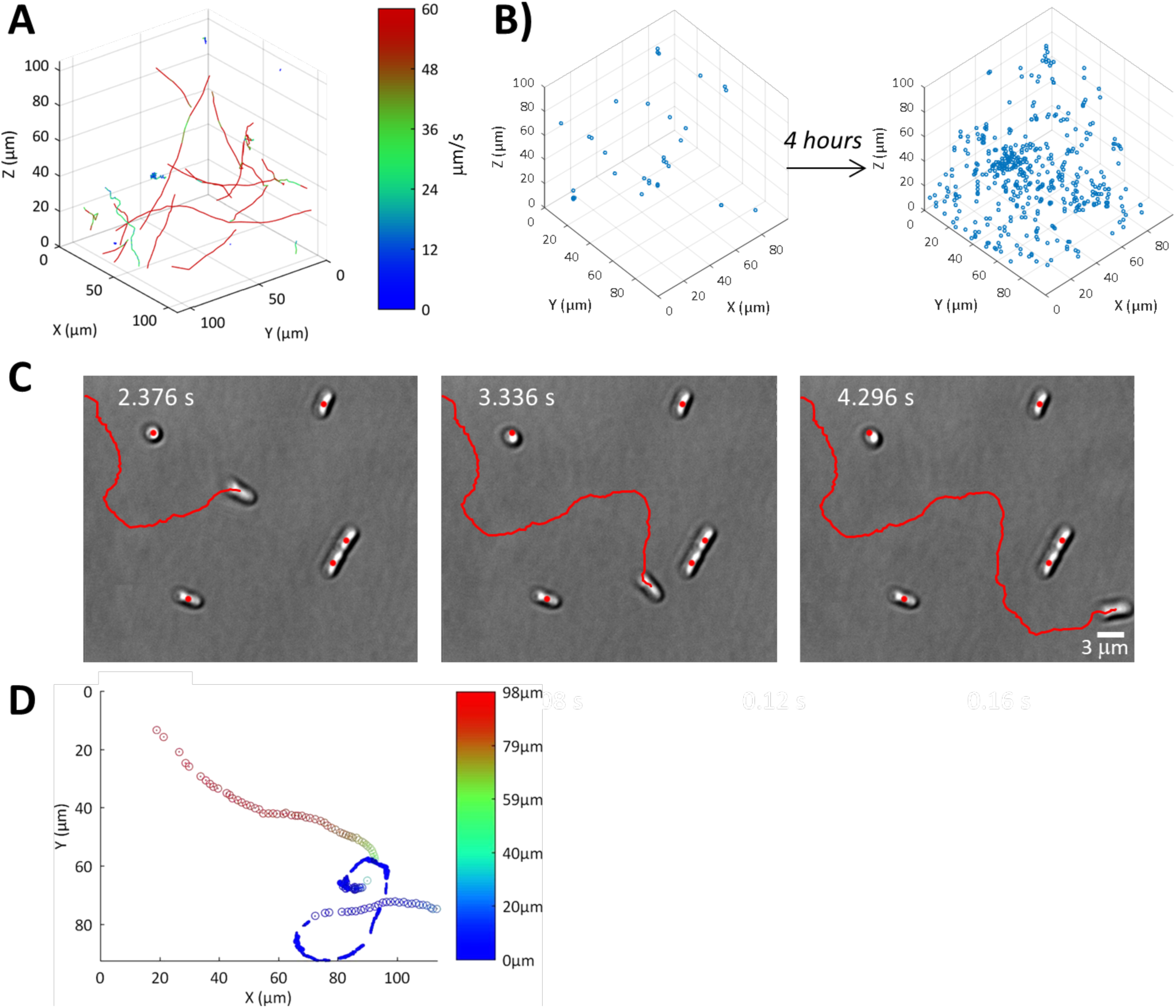
2D and 3D imaging of *P. aeruginosa* wild-type bacterial trajectories. (A) DHM isometric view of bacterial trajectories for one continuous 20 s acquisition at 41.7 Hz. Trajectories have been colored by the instantaneous speed of the track according to the intensity scale provided. (B) DIC Z-stack of instantaneous bacterial 3D positions at seeding conditions and after 4 h incubation acquired using the remote image relay illustrating the high cell density that can be located and tracked using this method. (C) DIC image sequence of *P. aeruginosa* cells swimming at the glass surface with 960 ms between images shown (complete image sequence is available in the supplementary material Movie S1). The trajectory is shown in red. Of note is the change in bacterial orientation that matched the change in direction during the journey. The image sequence also contains examples of long-axis (side) and polar (end) attached cells. The centre of mass of each cell at time = 2.376 s is indicated as a red dot in each frame, the centre of mass of side attached cells remained unchanged whereas a change in centre of mass location was observed for the end attached cell. (D) Example of a bacterial trajectory where a cell approached the surface from the bulk. The Z-position of the bacteria is coloured according to the intensity scale. The centre of mass as determined by DHM measurements is shown as a circle. When the bacteria came close enough to the surface the dimension of the bacteria was plotted as determined by DIC. All bacterial positions have been overlaid.

**FIG. 3.**
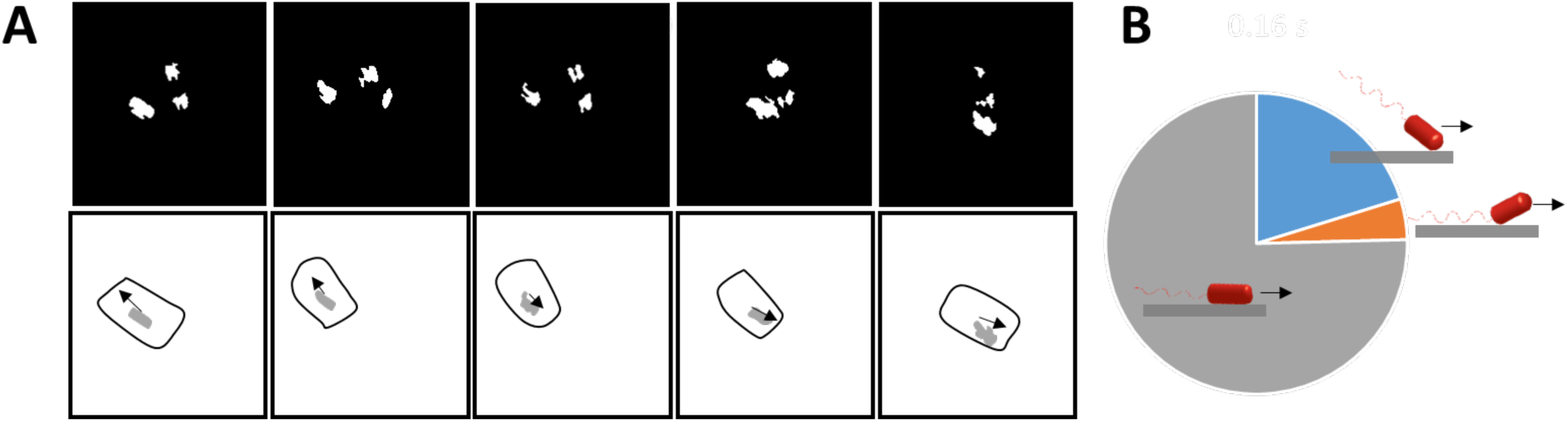
Interlaced capture of near-surface swimming bacteria with TIRM and DHM. (A) TIRM image sequence showing a direction reversal. Time interval of 40 ms between each displayed frame. The complete image sequence is available in the supplementary material (Movie S2). Images shown have been binarised and objects with an area less than 100 pixels, deemed to be noise, were removed. To aid in visualisation the outline (determined from DHM) of the moving cell is shown beneath the corresponding composite image along with the smoothed area of the cell imaged by TIRM (grey). The displacement vector of the centre of mass of the cell (determined from DHM) is also shown as an arrow. (B) Distribution of the position of the centre of mass of the TIRM spot within the cell body for 9 cells observed to be swimming parallel to the surface over a total of 183 frames, either at the front pole (blue), middle (grey) or rear pole (orange) of the cell (shown schematically where the arrows indicate the direction of travel of the bacterium). The front and rear of the cell were determined from the direction the bacterial cell was travelling subsequent to an image being taken. The TIRM spot was assigned as middle if it was less than 1 micron from the centre of mass of the cell as determined by DHM.

Single cell bacterial trajectories were determined using DHM (Fig. 2A) at cell densities of up to 1.7 × 10^4^ cells/μL, but above this, overlapping interference patterns prevented digital holographic reconstruction. To measure the 3D distribution of cells at higher populations we used the remote image relay to acquire a series of DIC images separated in the z-axis, referred to as *z-stacks*. Analysis of the z-stack data enabled the position of each bacterium to be determined at a population density of 3.7 × 10^5^ cells/μL, greater than an order of magnitude improvement over DHM, with a maximum population density within any one image of 8.0 × 10^5^ cells/μL. This enabled the growth of a *P. aeruginosa* cell culture to be followed over 4 h (Fig. 2B). Objects that were vertically contiguous for adjacent frames were attributed to a single bacterium and the position of each bacterial cell was determined as the centre of mass calculated from all pixels taken for combined objects. We employed a z-spacing of 1 μm in order to ensure that we did not omit bacterial cells when imaging (typical cell dimensions = 2 x 5 μm, depth of field at 60× magnification NA = 1.4 ≈ 0.6 μm). Over a z-range of 100 µm and a maximum imaging rate of 100 Hz, this meant that the maximum acquisition rate using the DIC z-stack was 1 Hz. Although useful for determining cell population density and distribution, the lower acquisition rate of this approach compared to DHM made it impractical for tracking fast swimming bacteria since, for accurate tracking, bacteria should not travel further than their dimensions per frame, thus, limiting this approach to cell movement slower than 5 μm/s. As such, DIC z-stacks were not used for tracking in this study but rather to assess cell distribution at high cell density.

### Characterization of bacterial trajectories

A number of different bacterial swimming trajectories were readily visually discernible within *P. aeruginosa* populations (Fig 2A **and** Fig 4). Relatively straight trajectories with cells travelling at high speeds are referred to as *runs* after the convention of Berg et al (5, 6) (Fig. 4A). Cells swimming parallel to the surface, were termed *near surface runs* (Fig. 4B), a phenomenon that has been assigned to various mechanisms including hydrodynamic interactions, Brownian motion and surface contact (7, 27, 35). Visual inspection of the tracks indicated that near surface runs had a greater incidence of directional change events than those observed in the bulk where trajectories were straighter for longer. Moreover, we observed circular trajectories where bacteria were travelling at the surface consistent with previous observations (Fig. 4B); curved near surface runs have previously been attributed to flagella-mediated surface interactions (13).

**FIG. 4.**
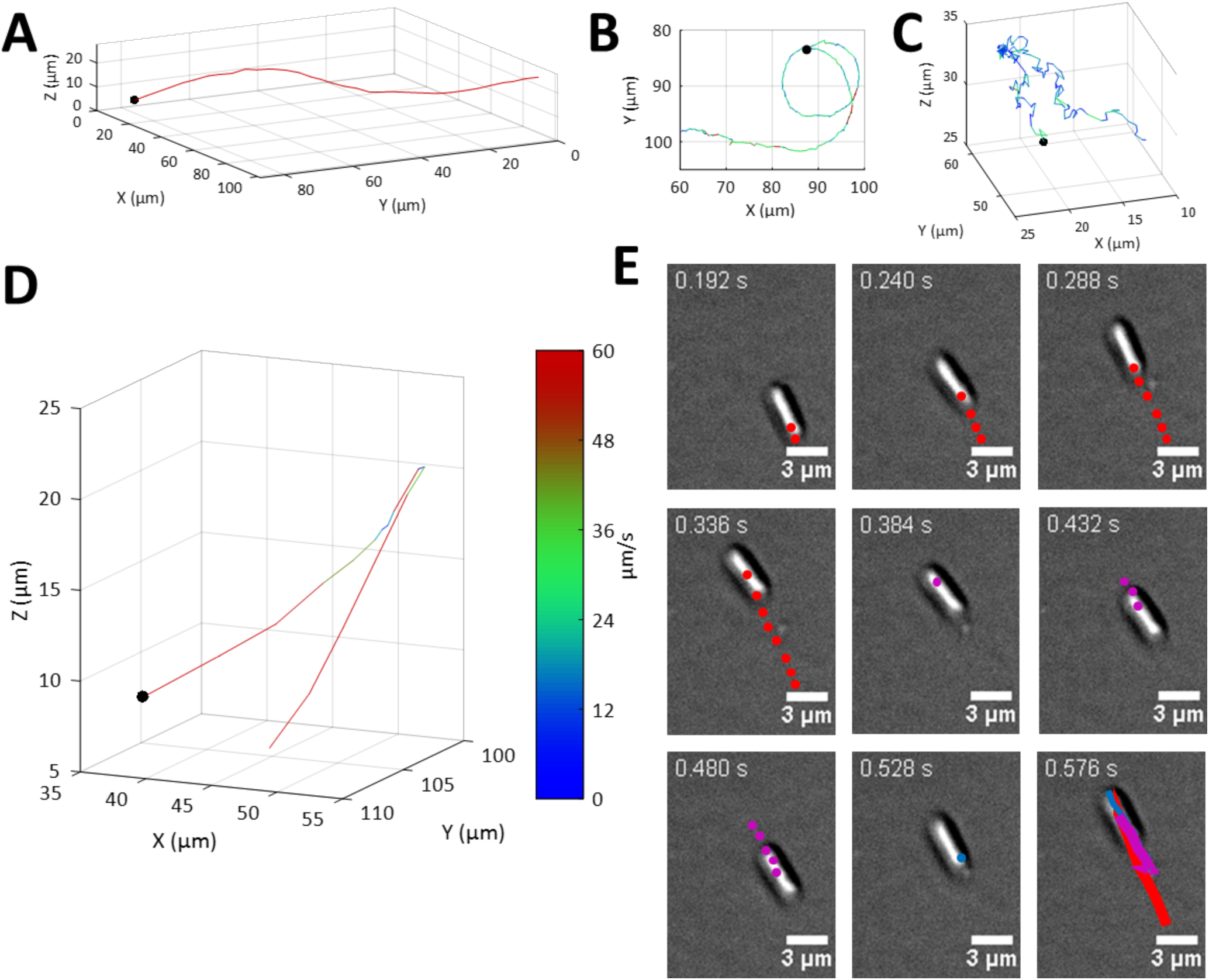
Characterization of bacterial swimming trajectories. (A-C) A number of different trajectory phenotypes were observed including: (A) run trajectory characterized by relatively long stretches of uninterrupted straight motion (track duration = 1.536 s), (B) near surface run with increased curvature (track duration = 4.872 s) and, (C) oscillating trajectories characterized by frequent changes in direction (track duration = 9.840 s). A black dot marks the beginning of each track. The track is colored by the instantaneous speed according to the intensity scale shown in (D). (D) Example of bacterial reversal directional change as observed by DHM within the bulk. The start of the trajectory is indicated by a black dot. The track is colored by the instantaneous speed according to the intensity scale. Track duration = 0.720 s. (E) Image sequence of two reversal events measured at the surface by DIC. The time between images is 48 ms. The center of mass of the cell from previous frames is shown as colored dots. The trajectory consists of the sequence forward-reverse-forward, and each segment has been colored red-magenta-blue, respectively. The entire trajectory is shown as colored lines on the final image shown. The two reversal events were observed around time points 0.336 s and 0.480 s whereupon the cell was observed to pause for a frame such that the center of mass of the cell remained unchanged between two frames.

Using TIRM, we showed that the bodies of near surface migrating bacteria were swimming parallel to the surface at a distance of less than 200 nm (Fig. 3A), a separation substantially shorter than the typical length of a flagellum (≈ 5 μm, Fig. S2A) (4). Through interlaced capture of DHM and TIRM imaging, the centre of mass of the bacteria (as determined by DHM) and the centre of mass of the contact point (as determined by TIRM) were compared (Fig. 3A). The majority of cells (75 %) exhibited a similar centre of mass and contact point, consistent with cells swimming parallel to the surface. In some cases, bacteria swam with either the front pole (20 %) or the rear pole (4 %) of the cell closer to the surface, defined by its direction of travel, consistent with the cell swimming front pole up or down (Fig. 3B). The measurements presented assume that the bacteria are in contact with the surface. This assumption is likely not true in all cases such that the number of cells swimming front pole up or down may be underestimated. This observation of monotrichously flagellated *P. aeruginosa* differs from observations of the swimming of the peritrichously flagellated *E. coli*, where all cells were reported to swim with the front pole down to the surface as estimated using 3 colour DHM (8).

Away from the solid-liquid interface, an alternative class of swimming trajectory was observed, where the movement of the bacteria was slower than the *near surface runs*, and characterized by frequent reversals of swimming direction, a phenomenon assumed to be caused by reversals of flagella rotation (Fig. 4C), termed *oscillating* after the classification applied by Rosenhahn et al (44). These oscillating bacterial cells were travelling at a reduced speeds of 12 ± 5 μm/s, substantially slower than the cells exhibiting run trajectories at 59 ± 4 μm/s. Notably, we did not observe *oscillating* cells at the surface, likely because the constraints of the proximal surface restricted the directional change of the cells upon flagella reversal. *Near surface runs* which we term *rambling* were slower and exhibited more frequent changes in direction than bulk *runs*. This rambling motion has not previously been reported. *Rambling* at the surface did not include the high frequency of directional reversals seen in the bulk (>90 degree) for oscillating cells.

Some cases were observed where *P. aeruginosa* trajectories were reminiscent of peritrichously flagellated *Escherichia coli* run-and-tumble patterns (5, 6), whereby a straight ‘run’ path was interspersed by a high frequency (2 Hz) of reversal events before the bacteria recommenced a run trajectory (Fig. S2B). The average reversal rate observed in the bulk population was 0.5 Hz. We observed no changes in the orientation of the bacterial body for surface reversal events (where the change in direction exceeded 120° - Fig. 4D) observed at the glass surface (Fig. 4E), consistent with these reversal events being caused by a change in flagellum rotational direction inducing a reversal of the bacterial direction of travel without reorientation of the bacterial body (37). To observe this more closely, we imaged cells using a third of the camera’s chip size enabling a higher acquisition rate of 333 Hz. At this higher frame rate we were still unable to observe any reorientation of the bacteria (**Movie S3**). Moreover, equipping the microscope with a fluorescence filter set enabled the use of total internal reflectance fluorescence (TIRF) imaging in order to visualize flagella through the use of fluorescent staining. Using this approach, flagella were readily observed at both the leading and trailing end of bacteria before and after surface reversal events (Fig. S2C, Movie S4), showing that *P. aeruginosa* cells are able to use their flagella to both push and pull, consistent with previous observations for monotrichous bacteria (12, 20, 32). We observed no statistical differences in the speed of bacterial trajectories before or after a reversal event (Fig. S2D).

Stationary cells at the surface were found attached either horizontally or vertically (long-axis or pole attached). Long-axis attached cells were identified from DIC images where the distinct rod-shape of the bacteria was observed and remained stationary throughout the duration of an image sequence with no change in the centre of mass of the cell (Fig. 2C). In contrast, pole-attached bacteria were observed to be smaller and circular in appearance and small deviations in the cell’s centre of mass were observed as the cell wobbled on a single attachment point (Fig. 2C). In some cases, polar attached cells were observed to be spinning due to the flagella being fixed to the surface whilst the bacteria body remained free and able to rotate, a phenomenon observed previously (14) (**Movie S5**).

### Exploring the respective contributions of the MotAB and MotCD stators to swimming motility

To exemplify the applications of the multimode microscope, we investigated the relative contributions of the MotAB and MotCD stators to swimming motility by comparing the trajectories of the *P. aeruginosa* wild type with the corresponding *motAB*, *motCD* and *motABCD* deletion mutants in order to assess cell behavior both within the bulk and at the surface. By tracking bacteria over a 2.4 s interval (Fig. 5), we first characterised the trajectories of each strain to determine whether the different forms of cellular movement apparent on visual inspection of cell tracking videos could be quantitatively categorized. As anticipated, the *motABCD* mutant was non-motile (15, 39), with all trajectories observed from this strain having a speed below 10 μm/s and a K_MSD_ below 1.5 (Fig. S3P). The average speed and K_MSD_ observed for this strain was 5.3 μm/s and 0.86, respectively. Therefore, the *motABCD* mutant was not analyzed further.

**FIG. 5.**
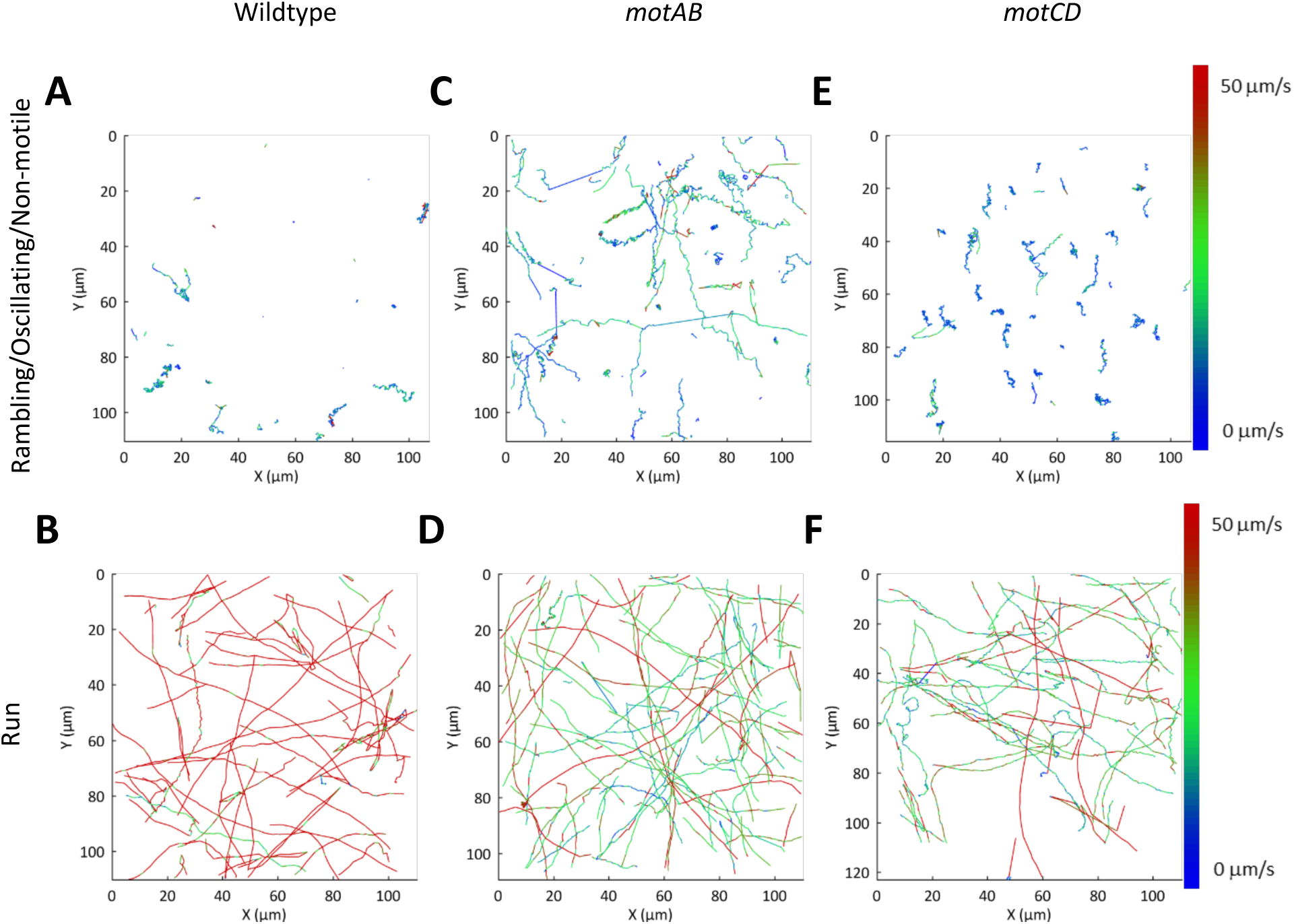
Top view of bacterial trajectories for *P. aeruginosa.* (A-B) wildtype, (C-D) *motAB*, and (E-F) *motCD* mutants, showing the trajectories coloured according to the speed (see intensity scale). Trajectories have been split into rambling/oscillating/non-motile (A,C,E) and run (B,D,F) trajectories.

Cell tracks were quantitatively characterized by instantaneous speed, total displacement, and their mean square displacement (MSD) over the time range (δt) of 50 to 1000 ms (Fig. S3A-I). The slope of the log plot (K_MSD_) was used to assess the directionality of a track, whereby a slope < 1 suggests constrained motion, a slope = 1 suggests Brownian diffusion and a slope > 1 suggests active motion (Fig. S3A-I) (11, 38, 48).

Considering the mean speed and K_MSD_ together, it was evident that two trajectory types were apparent in the *P .aeruginosa* wildtype population at both the surface and in the bulk (Fig. S3J-O). A cluster of tracks was observed for cells with a mean velocity above 30 μm/s and a K_MSD_ above 1.5, with the remaining trajectories having a mean velocity below 20 μm/s or a K_MSD_ below 1. A similar grouping of trajectories was observed for the *motAB* and *motCD* mutants, although in both cases the separation of the two groups was less evident due to a reduction in the velocity observed for the directionally swimming bacteria (Fig. S3K-L). Trajectories were separated into two classes, the first included bacterial tracks with a high velocity (> 25 μm/s) and a high K_MSD_ (>1.5), whilst the remaining were grouped into the second. Plots of the resulting tracks divided into the two classes are shown in Fig. 5. It was evident that based upon the selection criteria described, the trajectories separated into two clearly distinct motility types; in the first, the bacteria moved over large distances in a highly directional manner which have previously been described as *runs,* (Fig. 4A) and in the second, frequent reversals in direction were evident, termed *oscillating* (Fig 4C). Trajectories that remained at the surface were assigned as *rambling* as opposed to *oscillating*. We assigned runs that remained near the surface (within the focal plane of DIC images) for greater than 1 s as *near surface runs* (Fig. 4B). We were thus able to automatically assign the trajectory type and exclude the possibility of operator bias. The different bacterial trajectories were individually assessed to determine frequency, average speed and K_MSD_ (Table 1).

Using the analysis described, trajectory types were compared at the surface and within the bulk. The bulk bacterial wild-type trajectories were predominantly (64 %) runs, at an average speed of 59 ± 4 μm/s (mean ± standard deviation). In comparison both *motAB* and *motCD* mutants were predominately oscillating (≈54%) and the average run speed was significantly (p<0.0001) reduced in both cases to 29 ± 9 μm/s and 45 ± 18 μm/s, respectively (Fig. 6A; Table 1). This average measure of single-cell bacterial swimming contrasts with conventional measurements of population swimming speeds in low-viscosity agar whereby under these conditions no change in speed was observed for *P. aeruginosa motAB* and *motCD* mutants (40). In this prior study different classes of motility were not taken into account.

**FIG. 6.**
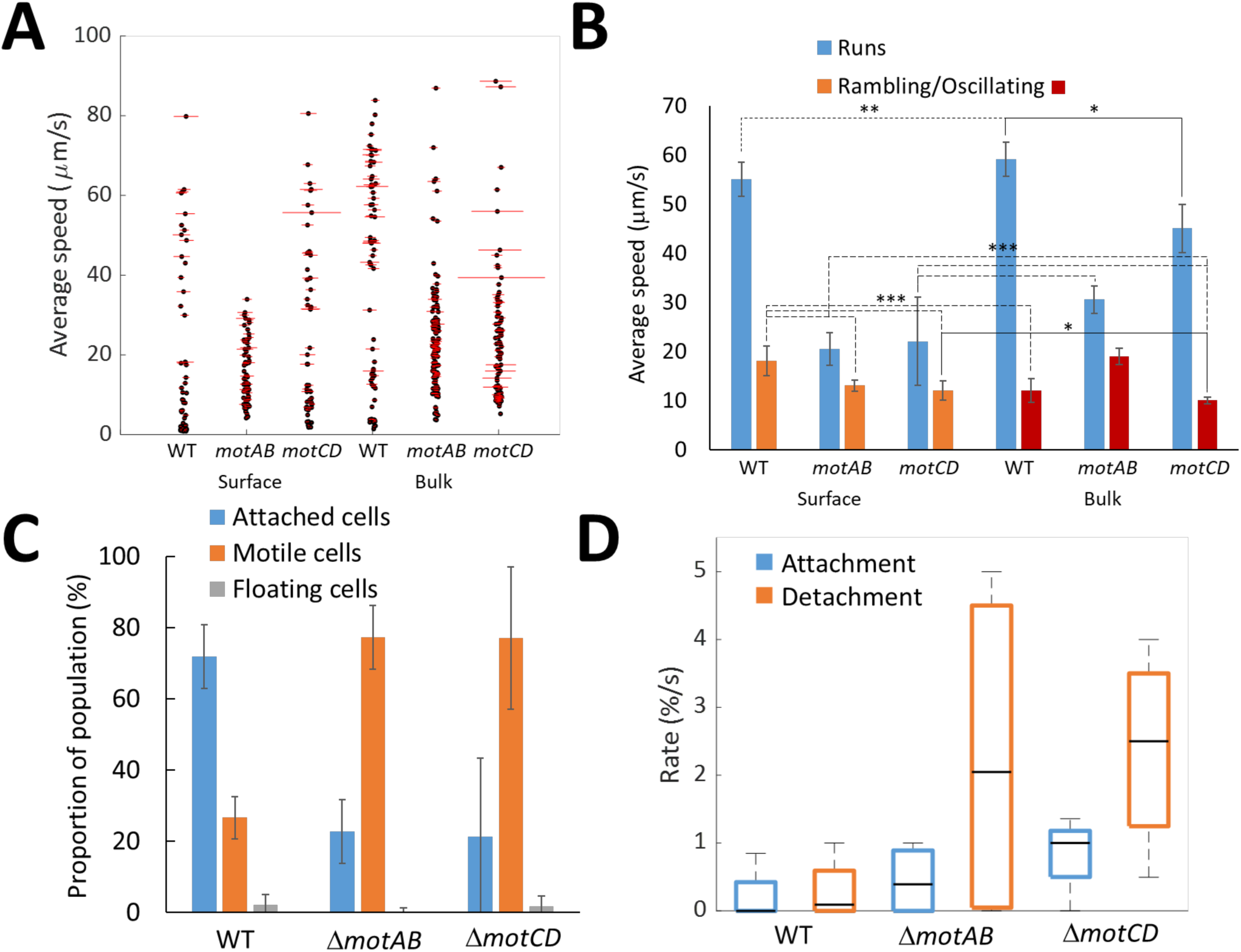
Comparison of the surface (DIC) and bulk (DHM) motility of *P. aeruginosa* wildtype and *motAB* and *motCD* mutants. (A) Comparison of the bulk and surface runs for speed. Each black dot is the measured average speed for a trajectory whilst the length of the associated red line is proportional to the standard deviation of the speed measured between frames over the trajectory length. (B) Comparison of the average speed measured for the run 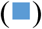, oscillating 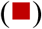 and rambling 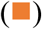 trajectories of *P. aeruginosa* wildtype and *motAB* and *motCD* mutants. The mean value is taken by averaging the speeds across 5 biological replicates. Error bars equal ± 95% confidence limits. Statistically significant differences of p<0.05 (*, solid line), p<0.01 (**, dotted line) and p<0.001 (***, dashed line) are shown. No difference or p<0.0001 differences are not shown. (C) Assessment of the attachment profile for each bacterial strain, reporting the total proportion of attached cells 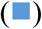, motile cells 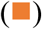 or floating cells 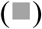. Error bars equal ± 1 standard deviation unit, N = 5. (D) Box and whisker plot for the attachment 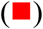 and detachment 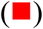 rates measured at the surface and reported as the % of non-attached cells attaching per s or the % of attached cells detaching per s. The central black line marks the median, the top and bottom of the box indicate the 75^th^ and 25^th^ percentile, respectively, whilst the whiskers extend to the maximum and minimum values, N = 5.

Increased variance in the speed within a single run trajectory was observed for the *motCD* mutants in the bulk (average standard deviation = 32 ± 36 μm/s compared with 18 ± 4 μm/s and 9 ± 1 μm/s for the wildtype and *motAB* strains, respectively. Fig. 6A, Table 1). Thus, removal of the *motCD* stator reduced the ability of the bacteria to swim at a constant speed.

The average speed of near surface runs was significantly (p<0.01) reduced compared to the average run speed in the bulk for all strains by 7 %, 39 % and 51 % for the wildtype, *motAB* and *motCD* mutants, respectively (Fig. 6A-B, Table 1). Thus, strains lacking either stator reduced their speed more significantly at the surface than the wildtype strain and, therefore, either reduced their power input or were unable to achieve the same speed for a certain power input within the fluid dynamics experienced at the liquid-solid interface (43, 50). This demonstrates a difference in the response to the surface between the stator mutants and the WT, consistent with a key role for the stators in responding to the surface environment as bacterial cells shift from the bulk environment to the near surface environment. Here, observation of the differential behaviour of the bacteria at the surface and bulk was achieved through the interlaced capture of cells both within the bulk and at the surface. The directionality of the run trajectories for all strains (including the wildtype) was unaltered either in the bulk or at the surface, with a K_MSD_ of 1.8 to 1.9 observed in all cases (Table 1).

In an oscillating trajectory the bacteria rapidly changed direction (Fig. 4C) as a result of the reversal of bacterial flagella rotation. (Fig. 4E). Both the wildtype and *motCD* mutant had high oscillating frequencies of 3.1 ± 1.0 Hz and 2.6 ± 0.6 Hz, respectively, whereas the *motAB* mutant exhibited a reduced oscillating frequency of 1.5 ± 1.0 Hz (significant at p<0.05) (Table 1). The average instantaneous speed of the oscillating *motAB* mutant was also higher (19 ± 5 μm/s) than the two other strains (≈11 μm/s; Table 1. As oscillating trajectories were observed for both mutants, either stator appears to be sufficient to support this type of trajectory. However, the altered phenotype of the *motAB* mutant suggests a possible role for MotAB in controlling the oscillatory trajectory. The directionality of the oscillating trajectories over a δt of 50 to 1000 ms also varied between the two mutants. The K_MSD_ values of 1.2-1.6 were lower than the run trajectories, thus this trajectory type allow the bacteria to move in a more diffusive manner. A statistically significant increase in K_MSD_ was observed for the *motAB* mutant compared with both the wildtype (p=0.04) and *motCD* (p=0.001) that is consistent with the oscillating trajectory requiring a contribution from MotAB.

A statistically significant (p<0.001) reduction in average speed was observed between the run and rambling trajectories at the surface for the wildtype and *motAB* strains, but not for the *motCD* mutant due to the high variability in the speed of surface runs for different trajectories observed for this strain (Table 1). The reduction in average speed from 22 to 12 µm/s for surface runs and rambling, respectively, for the *motCD* mutant was consistent with the other strains. For all strains a statistically significant (p<0.05) reduction in directionality from a K_MSD_ of 1.8-1.9 to 1.5-1.7 was observed. No significant difference in the K_MSD_ values for the rambling trajectories of the different strains was observed.

In addition, surface attachment, detachment and reversals for the *mot* mutants were also compared. After 5 min a larger proportion (72 %) of the wildtype cells were attached to the surface compared with either of the *mot* mutants (≈22 %) (Table 1). Of the attached cells, variance was also observed in the ratio of pole-attached to long axis attached cells, with values of 0.42, 2.7 and 1.2 observed for the wildtype, *motAB* and *motCD* mutants respectively (Table 1). Switching from long axis to polar attachment has previously been associated with cell surface departure (13). Consistent with this, although no statistically significant differences were observed between the attachment rates, the detachment rates for both mutants was higher (≈ 2.3 % of attached cells per second) than the wildtype (0.2 % of attached cells per s; (Table 1). Path deviations greater than 120 degrees, classified as reversals, were observed very infrequently for *runs* or *rambling* tracks. The highest reversal rates were observed for the wildtype bacteria (0.5 ± 0.1 Hz) whilst both *mot* mutants had similar reversal rates of 0.1 ± 0.1 Hz. The reversal rates for all strains increased when the bacteria were involved in near surface runs. Whereas for both wild type and the *motAB* mutant, reversal rates increased 2-fold upon exposure to the surface, the *motCD* mutant increased reversals by 6-fold (Table 1). This result contrasts with reversal rates observed previously for *P. aeruginosa* strain PA14 (39), where the reversal rates observed for the respective *motAB* and *motCD* mutants were 2-3 fold higher than the wildtype. This is likely due to different *P. aeruginosa* strain or culture conditions used, specifically the PA14 experiments were conducted using stationary phase bacteria grown in M63 medium supplemented with glucose and with the addition of 3% Ficoll to increase growth medium viscosity and so slow motility to enable imaging. The load responses of MotAB and MotCD stators have also been well studied (2) and the order of magnitude differences we observed in the reversal rates of the *motAB* and *motCD* mutants was likely a response to the variation in viscosity. Together these results suggest that the stators are involved in determining the outcome of *P. aeruginosa* cell interactions with a surface, consistent with previous observations (39). A few non-motile cells (<2 %) were observed for all motile strain populations. We also observed flagellar driven spinning for 10-13% of attached cells for all bacterial strains with the exception of the *motABCD* mutant where no spinning was observed, consistent with the flagellar dependent nature of this phenotype (14).

As the motility of log phase bacterial cells was investigated, we observed individual cells in various stages of cell division. To assess the influence of cell division on motility, we compared the trajectories of bacteria undergoing cell division identified via their elongated shapes. The near surface swimming behavior of dividing cells was unchanged compared with non-dividing cells, with no statistically significant differences observed between the speed or directionality of runs. However, the average speed of swimming, dividing cells was reduced from 40 ± 28 μm/s to 30 ± 17 μm/s, suggesting that during cell division, bacteria swim more slowly, likely as a result of the energetic costs of dividing, competition between the two daughter cell flagella that may not be fully formed, and the increased drag associated with a larger cell body.

To assess the role of the bacterial stators on bacterial surface accumulation, we observed the distributions of the *P. aeruginosa* wildtype and *mot* mutants after 2 h growth. Histograms of the bacterial distribution in the z-axis are shown in Fig. 7. A higher association of the wildtype and the *motAB* mutant with the two glass surfaces compared to the bulk was observed, both at the top and bottom (Fig. 7). For the wildtype bacteria the cell fraction (per μm) was more than 15× greater within 5 μm of the top and bottom surfaces compared with near the surface (5-20 μm) or within the middle of the sample chamber (20-50 μm). Assessment of the cell track classifications reveals that the surface accumulation was due both to near surface swimming and surface attachment. The surface accumulation observed for the *motAB* mutant was significant (p<0.05) but not as great as for the wildtype, with the cell fraction per μm at the top and bottom surfaces exceeding the middle of the sample chamber by 6-8 times. The cell fraction per μm was twice as great for the top and bottom surfaces for the *motCD* mutant compared with the middle of the chamber whilst for the *motABCD* mutant only the bottom surface showed a difference from the other regions. These differences, however, were not statistically significant (p>0.05). The minimal surface accumulation for the non-motile *motABCD* mutant was as anticipated given its lack of a flagellum. For the *motCD* mutant, the reduction of surface accumulation for this strain suggests a role for MotCD in surface sensing during the initial attachment stages. Moreover, the similar bulk and surface distributions of the motile *motCD* mutant suggests that the surface accumulation of bacteria cannot be explained by hydrodynamic or steric forces alone but may additionally be actively achieved by altered bacterial behavior when sensing a surface. No statistically significant differences were observed between the top and bottom surfaces for any of the bacterial strains suggesting that neither gravity nor buoyancy nor chemotaxis/aerotaxis are controlling factors in this experimental set-up (p>0.06).

**FIG. 7.**
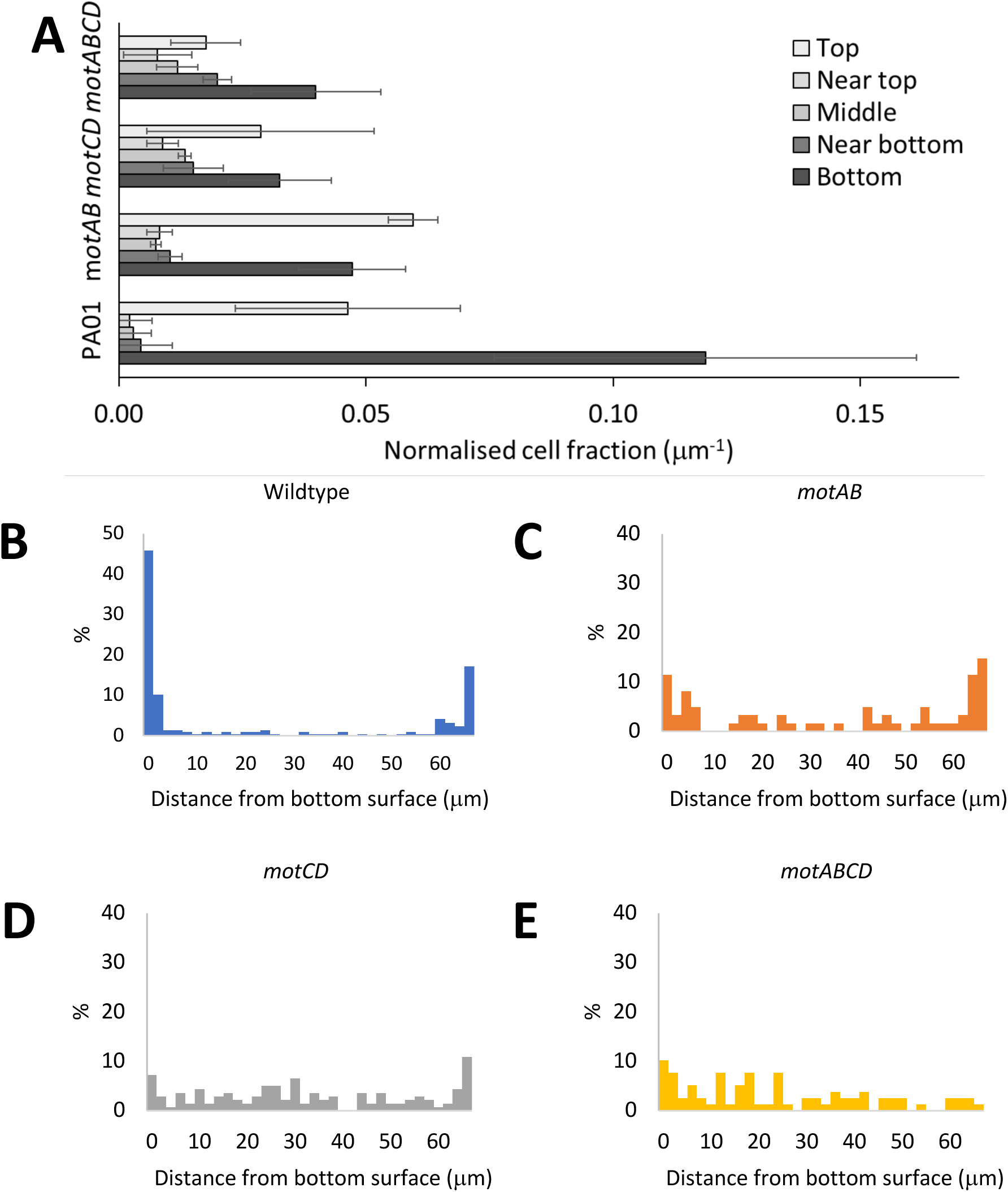
Investigation of surface accumulation of bacteria. (A) The normalized cell fraction (normalized to unit length in z axis) observed at varied z-regions within the sample chamber. Un-normalized data shown in B-E. Regions studied were bottom and top areas equal to 0-5 μm from the surface, near top and bottom equal to 5-20 μm from the surface, and the middle region. The average number of bacteria measured within the volume for each strain after the 5 min was wild type = 54, *motAB* = 21, *motCD* = 27, and *motABCD* = 15. Error bars equal ± 1 standard deviation unit from 3 biological replicates. Area imaged was 120 × 120 μm. (B-E) Fractions of bacterial population within different z-segments within a glass sandwich taken approximately 5 min after addition to chamber. *P. aeruginosa* (A) wild-type, (B) *motAB*, (C) *motCD* and (D) *motABCD* mutants. Data presented as a % of total number of cells observed in the z-stack.

## Discussion

*P. aeruginosa* adopts one of two major lifestyles, a free living motile or a surface-associated, sessile lifestyle (14, 29, 34). Understanding how bacteria make decisions to switch between such lifestyles is important for unlocking new strategies for the prevention and treatment of bacterial infections, especially biofilm-centred infections given their chronicity and resistance to host defenses and antibiotics. The key role of surfaces in bacterial lifestyle choices highlights the importance of developing methods that are able to capture quantitatively the differences in bacterial behavior within populations at the surface and in the bulk in order to provide new insights into bacterial surface sensing. To this end, we have developed a multi-mode microscope with dual surface/bulk imaging capabilities able to observe subtle differences in the bacterial trajectories of *P. aeruginosa* cells within the bulk and the near surface environment.

Characterization of bacterial trajectories revealed two types of motility that were consistent with previous studies (37, 44), and could be categorized by consideration of speed and directionality. Notably, although runs were observed both at the surface and in the bulk, oscillatory trajectories were observed only within the bulk whilst rambling trajectories were observed only at the surface. Deletion of either stator did not inhibit any trajectory type, demonstrating that a specific stator complex was not associated with a specific trajectory type. However, the oscillatory frequency for the *motAB* mutant was reduced compared to the WT, suggesting a role for this stator in regulating this particular trajectory type. It was not clear what caused the bacteria to exhibit a particular trajectory type, however, the lower speed and decreased directionality of the rambling and oscillatory trajectory types suggest an explorative behavior in contrast to runs, which allow bacteria to travel more rapidly over larger differences. Adopting the rambling or oscillatory trajectories also enable bacterial cells to alter direction in a similar manner to the tumbling action observed for the peritrichous *E. coli* (5).

A greater distribution of average speeds was noted for cells within the *motCD* mutant population as a consequence of the greater variability in the speed of an individual cell compared with the WT or *motAB* mutant. Previous studies have demonstrated that bacteria are able to control their torque through regulating the turnover of stator units (9, 26). The variability in the speed of the *motCD* mutants as compared with the WT and *motAB* mutants suggests a greater variability in the number of stator units within the flagella complex across the population.

Whereas the WT strain swimming speed was statistically indistinguishable between bacteria swimming at the surface and those within the bulk, the *motAB* and *motCD* mutants swimming speed at the surface was significantly (p<0.0001) slower than cells within the bulk. It is possible that either the WT is able to adjust to the altered physico-chemical near-surface environment or to obstruction of flagella rotation by the surface due to the exchange of the two stator complexes to modulate flagella torque. Stator exchange has been proposed to act as a signaling mechanism in concert with c-di-GMP within *P. aeruginosa* (1, 3, 24). The absence of either stator within the *motAB* and *motCD* mutants removes the option for stator exchange, preventing the cells from being able to either sense or respond to the near surface environment using this mechanism.

The involvement of both bacterial appendages, flagella and pili, in bacterial swarming, biofilm formation and, ultimately, surface sensing has been reported (3, 23, 25, 29, 33). In *P. aeruginosa,* flagella signaling involves c-di-GMP and the diguanylate cyclase SadC (3), whilst pili, particularly TFP, are able to mechanically interact with a surface through successive pili extension and retraction that enables signal transduction through the Chp system to regulate cyclic adenosine monophosphate (c-AMP) levels (31). Whilst the mechanosensing role of pili for regulating the behavior of surface associated bacteria is clear, our results, and those of others (25) also support a role for the flagellum in surface sensing.

Here we have also shown that both the *motAB* and *motCD* mutant populations included a higher proportion of detached cells, and that attached cells were more likely to detach, further demonstrating the differential surface response of *P. aeruginosa* cause by the deletion of either of the flagella stators. This is likely caused by the inability of the bacteria to sense the surface effectively and so they fail to trigger the upregulation of c-di-GMP and pili required for cells to become irreversibly surface attached leading to biofilm formation.

The similar behaviors of both *mot* mutants suggest that both stators are involved in the cell signaling, potentially through stator exchange. The absence of surface accumulation noted only for the *motCD* mutant however, indicates a greater role for the MotCD stator in surface sensing and subsequent biofilm formation.

In summary, we have developed a multimode 2/3D microscope that combines DIC, DHM, TIRM and TIRF imaging modes to achieve simultaneous bulk and surface label-free imaging of single cells in a motile bacterial population. The microscope was used to investigate the role of the *P. aeruginosa motAB* and *motCD* flagella stators on motility and surface interactions, enabling observations of altered phenotypic behavior of cells located within the bulk or at the surface.

## Acknowledgments

This work was supported via Wellcome Trust joint senior investigator awards (Grant Nos. 103882 and 103884). Dr. A. L. Hook would also like to thank the University of Nottingham for funding via his awarded Nottingham Research Fellowship. The authors declare no competing conflicts of interest. Assistance from Andrew Woodward with Matlab scripts is kindly acknowledged.

## Supplementary Material

**Supplementary movies 1-5 are available for viewing.**

MOVIE S1 – DIC image sequence of *P. aeruginosa* cells swimming at the glass surface showing rambling cells, and example of cells side or end attached.

MOVIE S2 – Example of simultaneous capture of TIRM (left) and DHM (right), showing an example of a *P. aeruginosa* cell swimming at the glass surface.

MOVIE S3 – High frame rate (333 Hz) DIC image capture of a *P. aeruginosa* cell swimming at the glass surface. Image has been binarised (2× the standard deviation of the background pixel intensity) to improve visualization. Due to short exposure time single pixel noise is visible.

MOVIE S4 – TIRF image sequence using a 488 nm laser excitation and showing *P. aeruginosa* wild-type arriving and then leaving the surface, demonstrating push and pull swimming behaviour. TIRF evanescent field thickness set to 300 nm. The *P. aeruginosa* WT was stained with AlexaFluor 488 to enable visualization of the flagella. See also Fig S3.

MOVIE S5 – DIC image sequence of *P. aeruginosa* spinning at the glass surface.

**FIG. S1.**
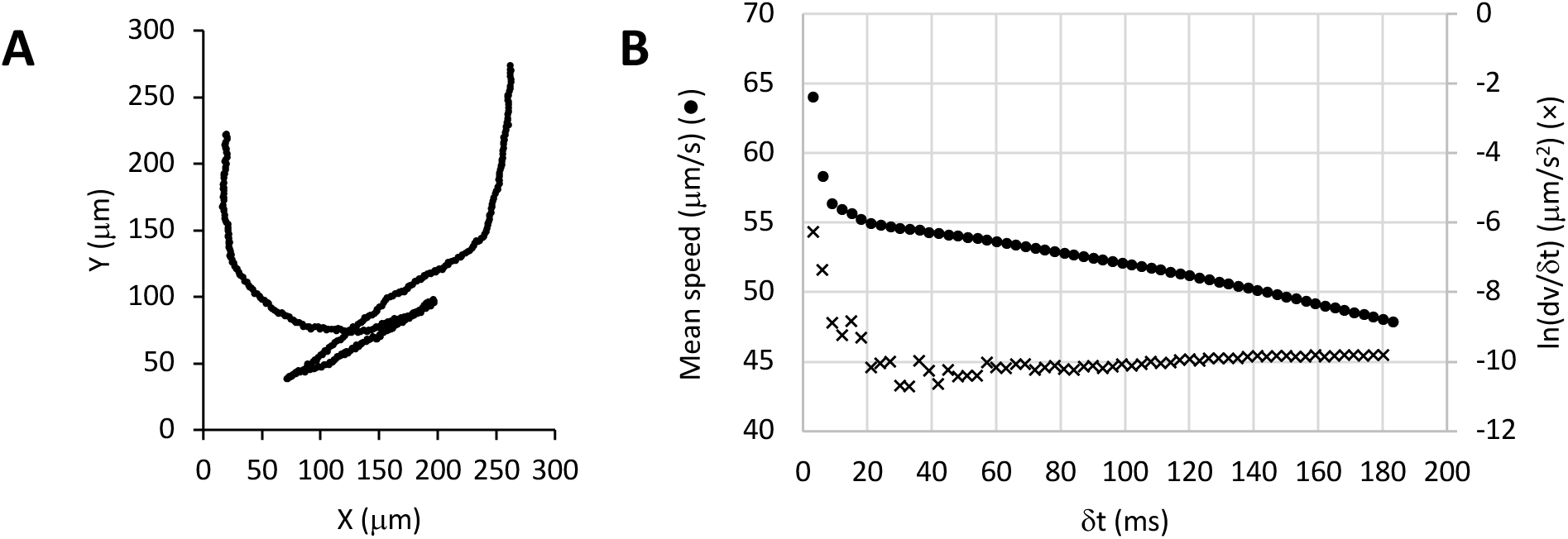
Assessment of capture rate with respect to sampling object speed. (A) X-Y coordinates of a bacterial trajectory taken with DIC. (B) The mean speed measured for a bacterial trajectory at high frame rate (333 Hz) when the speed was measured between different time intervals. The higher mean speed measured at shorter time intervals suggest that the bacteria experience fluctuating motion over a small time scale. The mean speed was observed to decrease to a minimum in the differential dν/δt within the region of 20-60 ms. Above 40 ms a further decrease in the measured mean speed was observed suggesting that at these time intervals the measurements were under-sampled with respect to the directional motion of the bacteria. Assuming the bacteria were a sphere of diameter = 3 µm, mass = 6.25 × 10^-13^ kg and a media viscosity of 0.6913 mPa.s at 37 ⁰C the Brownian motion frequency as determined by Stokes law = 3.12 × 10^4^ Hz, suggesting that Brownian motion is under-sampled at all imaging frequencies. See also supplementary movie 3 for full image sequence.

**FIG. S2.**
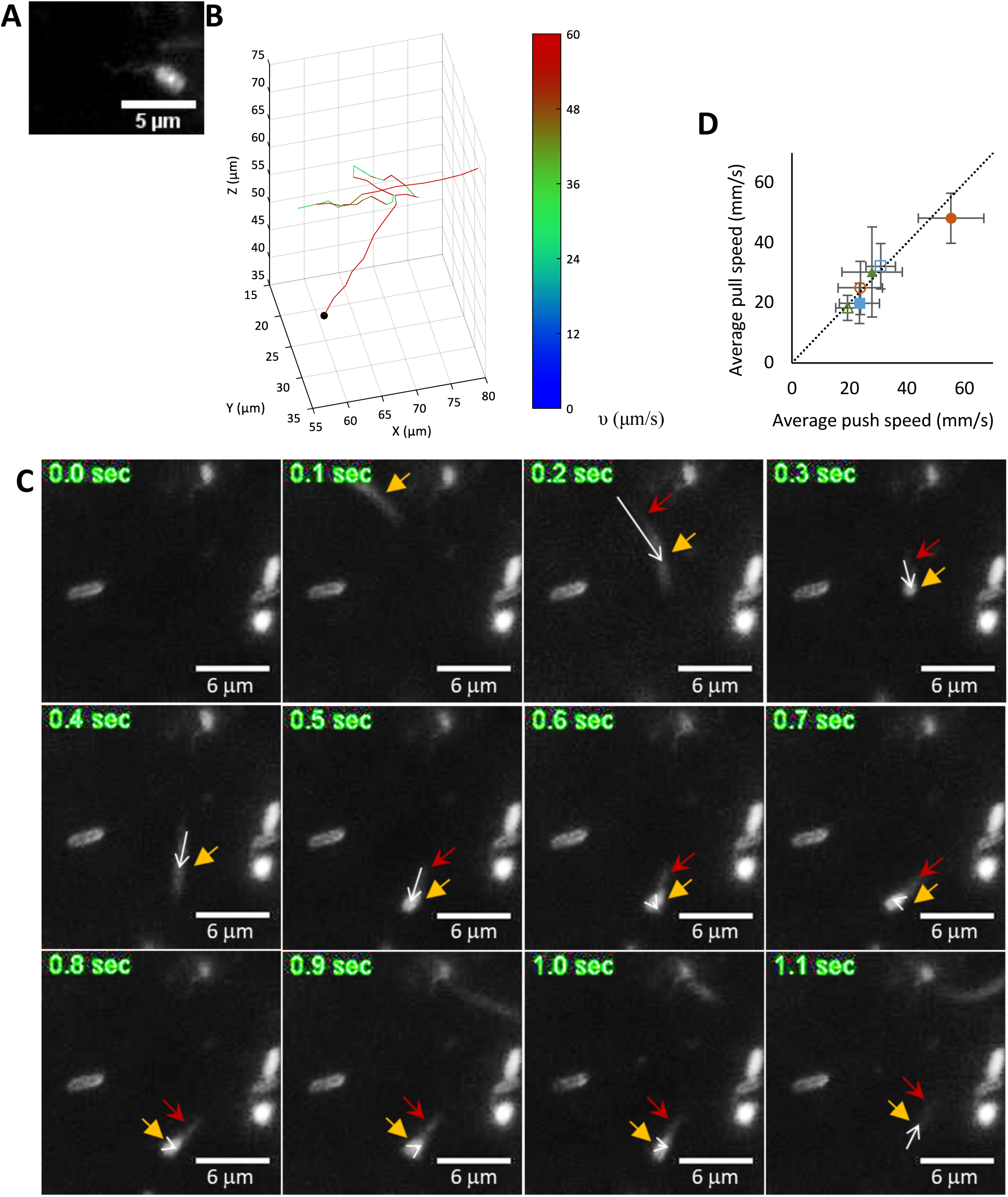
Investigation of reversing bacteria. (A) TIRF image of *P. aeruginosa* wild-type bacterium stained with AlexaFluor 488 to enable visualisation of the flagella. The coiled flagella length was measured to be 3.8 μm. (B) Example of run-reverse trajectory. The start of the trajectory is indicated by a black dot. The track is coloured by the instantaneous speed according to the intensity scale. (C) TIRF image sequence using a 488 nm laser excitation and showing *P. aeruginosa* wild-type arriving and then leaving the surface, demonstrating push and pull swimming behaviour. TIRF evanescent field thickness set to 300 nm. Image capture at 10 Hz using a 60×, NA=1.4 objective. The white arrows show the displacement vector between the centre of mass of the bacterial cell of interest between two frames. The yellow arrow identifies the centre of mass of the body and the red arrow shows the position of the flagellum (when visible) of the bacterial cell of interest. All other cells in the frame were stationary for the duration of the image sequence. (D) Comparison of push and pull speeds of *P. aeruginosa* wildtype (circles) with *motAB* (triangles) and *motCD* (squares) mutants in the bulk (closed) and at the surface (open). Error bars represent 95% confidence limits. See also Movie 4.

**FIG. S3.**
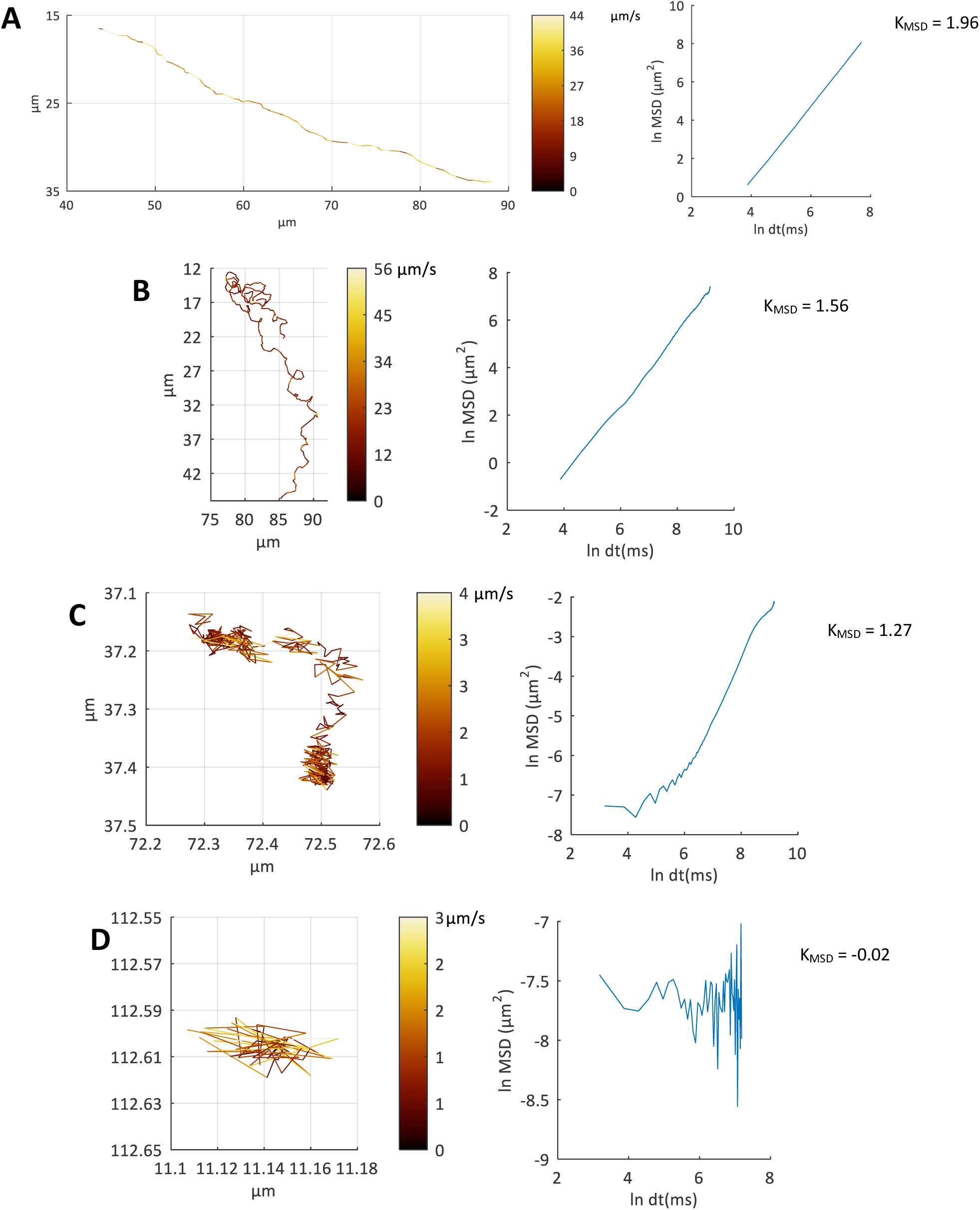
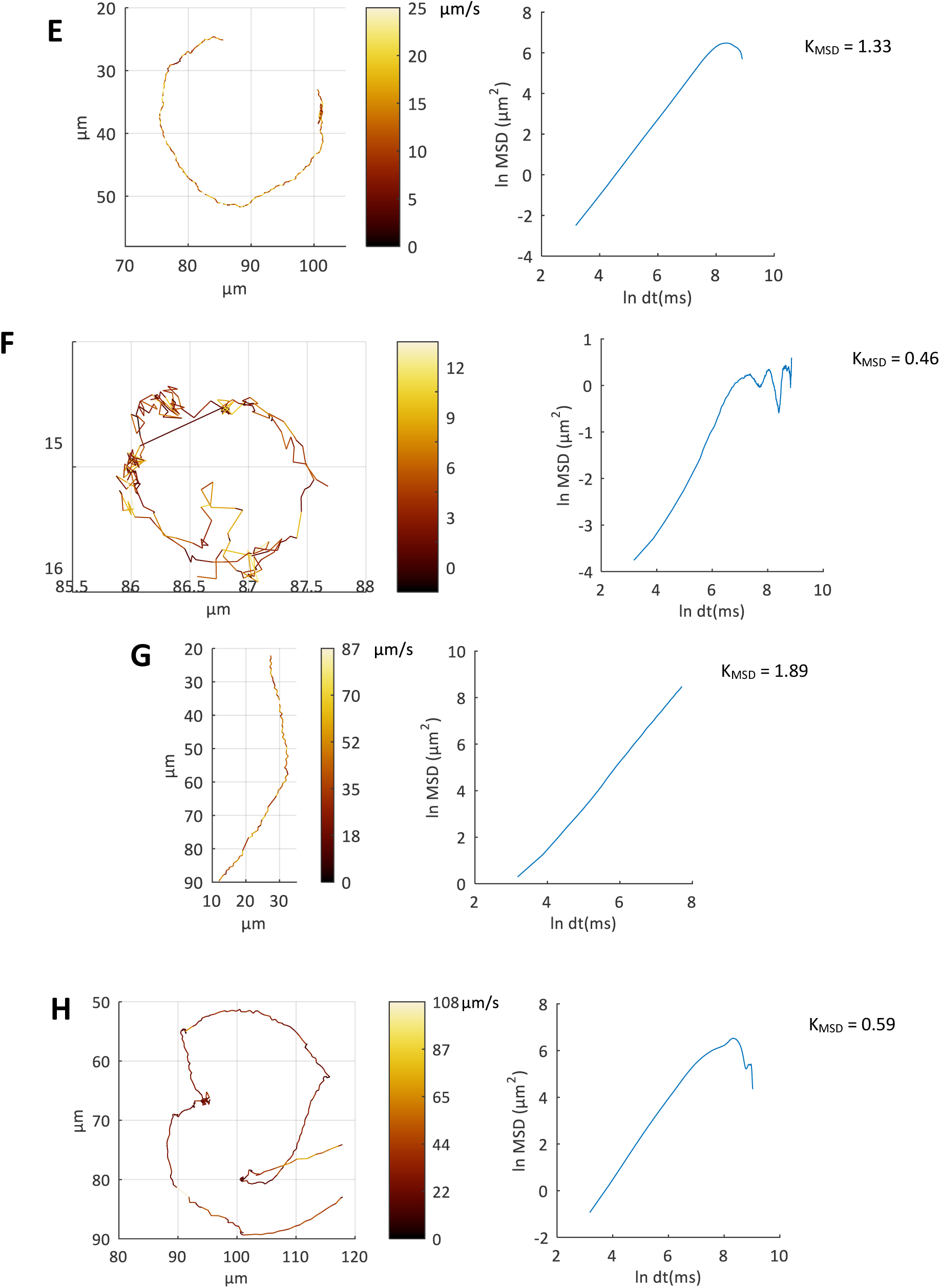
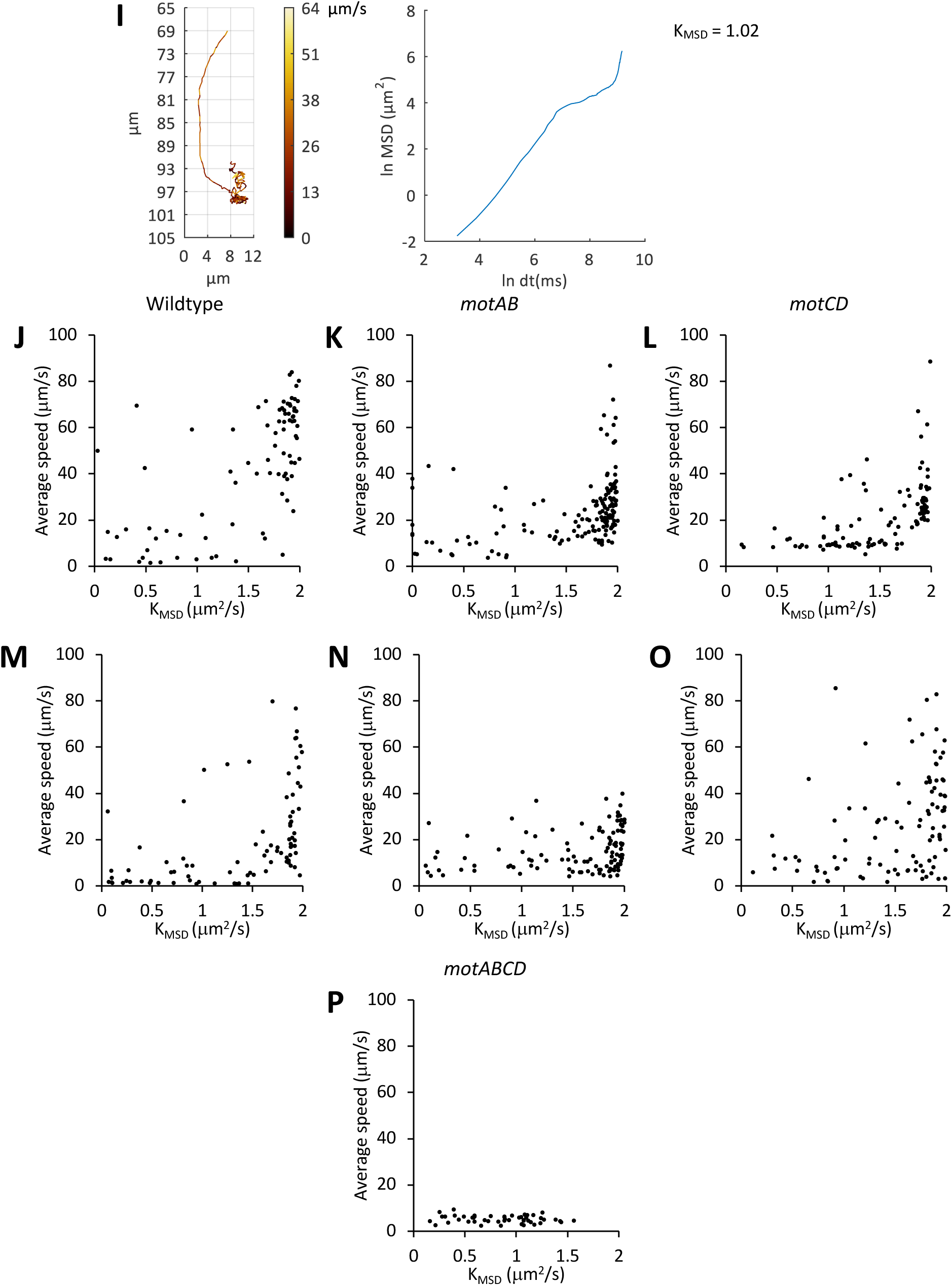
Exploration of the MSD and speed as characterisation of trajectories. (A-I) Examples of different track types, showing the X,Y position of the track (left) and the log-log plot of MSD at various time intervals dt (right). The X,Y coordinates of the track have been coloured according to the tracks instantaneous speed according to the intensity map provided in units of μm/s. The slope of the line of best fit through the log-log plot is shown (K_MSD_)^1^. (J-P) Plot of the mean speed (μm/s) and K_MSD_ for every track measured in the bulk (J-L) and at the surface (M-P) for (J, M) *P. aeruginosa* wildtype, (K, N) *motAB*, (L, O) *motCD* and (P) *motABCD* mutants from 4 biological replicates.

**TABLE S1.**
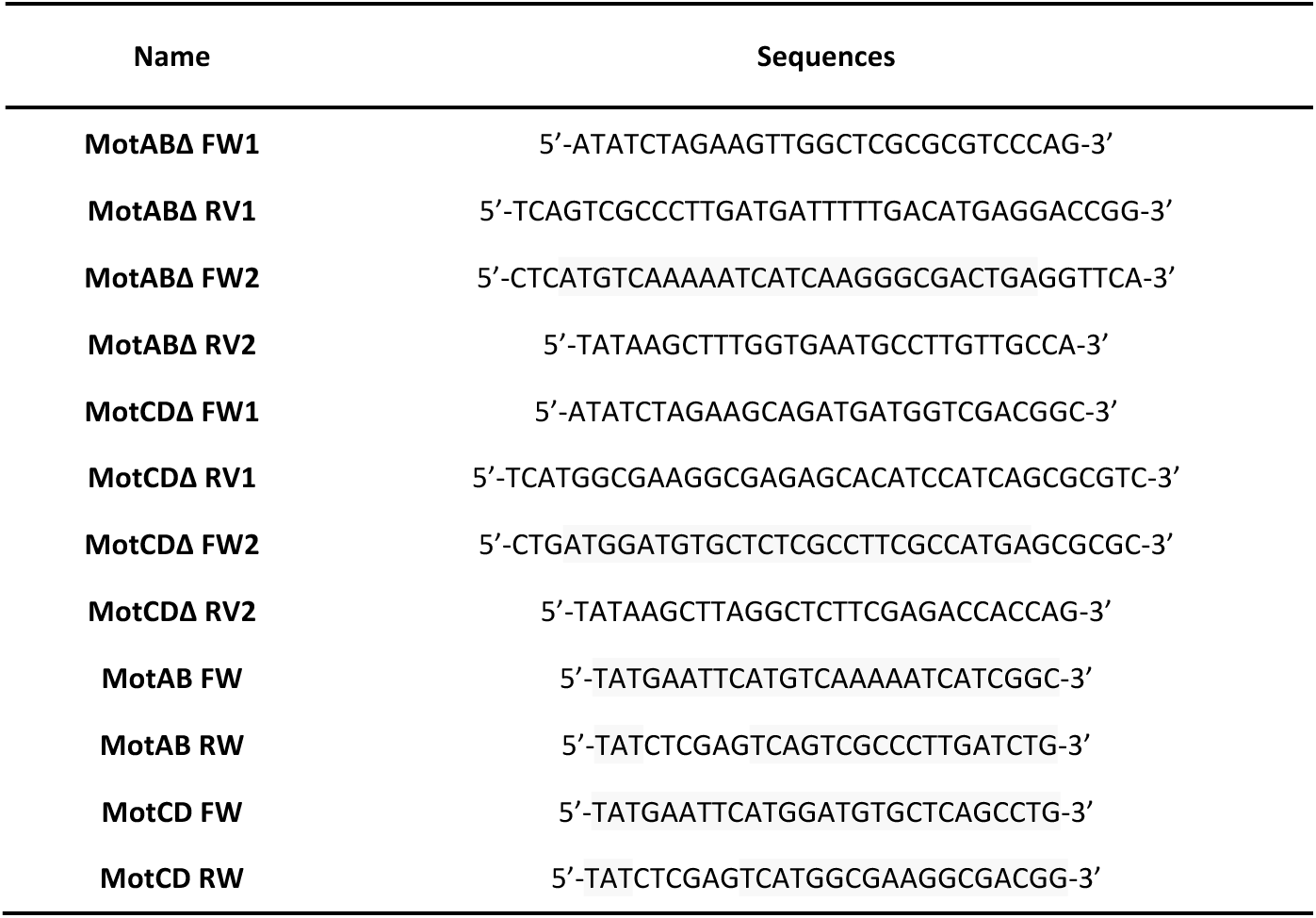
List of oligonucleotides used in this study for the construction of *motAB* and *motCD* in frame deletion muta<u>nts and for geneti</u>c complementation.

## Footnotes

1 K_MSD_ has widely been used to characterise the movement of particles, enabling directed, Brownian and confined movement to be defined. A number of different examples of bacterial tracks and the associated K_MSD_ measurements are shown in Fig. SI5. Tracks travelling directionally were observed to produce a linear relationship between the log of MSD over the log of time intervals with a slope approaching 2 (Fig. S5A). A track characterised by frequent oscillations had a reduced K_MSD_ of 1.56 (Fig. S5B), whilst the drifting of a non-motile bacterium had a K_MSD_ closer to 1 (Fig. S5C). A bacterium attached to the surface and therefore confined has a K_MSD_ of approximately 0 (Fig. S5D). A K_MSD_ below 2 was also observed for a bacterium travelling in circular trajectories (Fig. S5E) or spinning (Fig. S5F). In this case a linear relationship between the ln of MSD and ln of δt was observed up to the time interval associated with a complete oscillation, whereupon a plateau in the log MSD was observed at a position determined by the diameter of the curved path (Fig. S5F). In cases where the bacterium experienced oscillating movement over short time scales (fluctuating motion) and directional movement over larger times scales (mean motion) two slopes were observed on the log-log plot of MSD and δt (Fig. S5G), whereupon the two slopes were indicative of the directionality of the two movement types. As directional movement of the bacteria was of interest for comparing trajectories and the oscillating movement of the bacteria over short time frames was undersampled (Fig. S1) the K_MSD_ value was calculated over δt values of 50 to 1000 ms. Changes in the bacterial movement during a single track caused by reversal events or attachment or detachment events resulted in spurious measurements of K_MSD_ (Fig. S5H-I). For this reason tracks were split when attachment, detachment or reversal events were observed prior to K_MSD_ analysis.

## Notes

https://github.com/fishhooky/DHMTrack

